# Population sequencing data reveal a compendium of mutational processes in human germline

**DOI:** 10.1101/2020.01.10.893024

**Authors:** Vladimir B. Seplyarskiy, Ruslan A. Soldatov, Ryan J. McGinty, Jakob M. Goldmann, Ryan Hernandez, Kathleen Barnes, Adolfo Correa, Esteban G. Burchard, Patrick T. Ellinor, Stephen T. McGarvey, Braxton D. Mitchell, Vasan S. Ramachandran, Susan Redline, Edwin Silverman, Scott T. Weiss, Donna K. Arnett, John Blangero, Eric Boerwinkle, Jiang He, Courtney Montgomery, D.C. Rao, Jerome I. Rotter, Jennifer A Brody, Yii-Der Ida Chen, Lisa de las Fuentes, Chii-Min Hwu, Stephen S. Rich, Ani W. Manichaikul, Josyf C. Mychaleckyj, Nicholette D. Palmer, Jennifer A. Smith, Sharon L.R. Kardia, Patricia A. Peyser, Lawrence F. Bielak, Timothy D. O’Connor, Leslie S. Emery, NHLBI Trans-Omics for Precision Medicine (TOPMed) Consortium, TOPMed Population Genetics Working Group, Christian Gilissen, Wendy S.W. Wong, Peter V. Kharchenko, Shamil Sunyaev

**Affiliations:** Division of Genetics, Brigham and Women’s Hospital, Harvard Medical School, Boston, MA, USA; Department of Biomedical Informatics, Harvard Medical School, Boston, MA, USA; Department of Human Genetics, Radboud Institute for Molecular Life Sciences, Radboud University Medical Center, Nijmegen, the Netherlands; Quantitative Life Sciences, McGill University, Montreal, QC; Department of Medicine, University of Colorado Denver, Aurora, CO 80045, USA; Department of Medicine, University of Mississippi Medical Center, Jackson, MS; Department of Pediatrics, University of Mississippi Medical Center, Jackson, MS; Department of Population Health Science, University of Mississippi Medical Center, Jackson, MS; Department of Bioengineering and Therapeutic Sciences, University of California, San Francisco, CA; Department of Medicine, University of California, San Francisco, CA; Program in Medical and Population Genetics, The Broad Institute of MIT and Harvard, Cambridge, MA; International Health Institute, Brown University, Providence, RI; Department of Epidemiology, Brown University, Providence, RI; Department of Anthropology, Brown University, Providence, RI; Department of Medicine, University of Maryland School of Medicine, Baltimore, MD; Program for Personalized and Genomic Medicine, University of Maryland School of Medicine, Baltimore, MD; Geriatrics Research and Education Clinical Center, Baltimore Veterans Administration Medical Center, Baltimore, MD; Department of Medicine, Boston University School of Medicine, Boston, MA; Framingham Heart Study, Framingham, MA; Department of Medicine, Harvard Medical School, Boston, MA; Department of Medicine, Brigham and Women’s Hospital, Boston, MA; Channing Division of Network Medicine, Department of Medicine, Brigham and Women's Hospital, Boston, MA; Department of Epidemiology, University of Kentucky, Lexington, KY; Department of Human Genetics, University of Texas Rio Grande Valley School of Medicine, Brownsville, TX; South Texas Diabetes and Obesity Institute, University of Texas Rio Grande Valley School of Medicine, Brownsville, TX; University of Texas Health Science Center at Houston, Houston, TX; Baylor College of Medicine Human Genome Sequencing Center, Houston, TX; Department of Epidemiology, Tulane University, New Orleans, LA; Tulane University Translational Science Institute, Tulane University, New Orleans, LA; Division of Genomics and Data Science, Department of Arthritis and Clinical Immunology, Oklahoma Medical Research Foundation, Oklahoma City, OK; Division of Biostatistics, Washington University in St. Louis, St. Louis, MO; Department of Pediatrics, The Institute for Translational Genomics and Population Sciences, Los Angeles Biomedical Research Institute at Harbor-UCLA Medical Center, Torrance, CA; Department of Medicine, The Institute for Translational Genomics and Population Sciences, Los Angeles Biomedical Research Institute at Harbor-UCLA Medical Center, Torrance, CA; Cardiovascular Health Research Unit, Department of Medicine, University of Washington, Seattle, Washington, USA, 98115; Washington University, St. Louis, MO 63130; National Yang-Ming University School of Medicine, Taipei, Taiwan; Center for Public Health Genomics, University of Virginia, Charlottesville, VA, USA; University of Virginia Center for Public Health Genomics, Charlottesville, Virginia, USA; Department of Biochemistry, Wake Forest School of Medicine, Winston-Salem, NC; Department of Epidemiology, School of Public Health, University of Michigan, 1415 Washington Heights, Ann Arbor, MI 48109-2029; Survey Research Center, Institute for Social Research, University of Michigan 426 Thompson St, Room Ann Arbor, MI 48104; Institute for Genome Sciences, University of Maryland School of Medicine, Baltimore, MD; University of Maryland Marlene and Stewart Greenebaum Comprehensive Cancer Center, Baltimore, MD; University of Washington Department of Biostatistics, Seattle, WA 98195; Inova Translational Medicine Institute (ITMI), Inova Health Systems, Falls Church, VA, USA

## Abstract

Mechanistic processes underlying human germline mutations remain largely unknown. Variation in mutation rate and spectra along the genome is informative about the biological mechanisms. We statistically decompose this variation into separate processes using a blind source separation technique. The analysis of a large-scale whole genome sequencing dataset (TOPMed) reveals nine processes that explain the variation in mutation properties between loci. Seven of these processes lend themselves to a biological interpretation. One process is driven by bulky DNA lesions that resolve asymmetrically with respect to transcription and replication. Two processes independently track direction of replication fork and replication timing. We identify a mutagenic effect of active demethylation primarily acting in regulatory regions. We also demonstrate that a recently discovered mutagenic process specific to oocytes can be localized solely from population sequencing data. This process is spread across all chromosomes and is highly asymmetric with respect to the direction of transcription, suggesting a major role of DNA damage.

---

The superb accuracy of transmission of genetic information between generations is one of the most fascinating properties of life. Infrequent errors in this transmission lead to mutations that are the source of genetic variation which fuels evolution and causes genetic disease. The key importance of mutagenesis motivated decades of experimental research that revealed various modes of errors made by complex machineries of DNA replication and DNA repair (*1*–*3*). In spite of this effort, biochemical mechanisms primarily responsible for human germline mutation remain uncharacterized. Statistical analysis of massive whole genome sequencing datasets in light of the knowledge accumulated by experimental genetics and biochemistry offers a promising avenue of inquiry.

Studies of the origin of cancer somatic mutations have been propelled by the statistical analysis of “mutation signatures” in cancer genomic datasets and by mapping these signatures to known exposures to endogenous and exogenous mutagens (*4*–*6*). This analysis exploits the trinucleotide context-dependency of mutation rate. Differential exposure of tumors to mutagens serves as the main statistical instrument for the analysis. This approach is not directly transferable to studies of human germline mutation because there is no analog of the differential mutagen exposure, although some success was achieved by comparing human populations (*7*–*10*).

Here, we use variation along the genomic coordinate as the statistical instrument to decompose human germline mutagenesis into independent biochemical processes. Human mutation rate exhibits a modest but highly significant variation along the genome (*11*–*13*). Our model assumes that several mechanistic processes generate human germline mutations. These processes are characterized by types and context-dependency of nucleotide changes and vary in their relative intensities along the genome (Fig. 1A). Mutational signatures and the relative intensity of each process at each locus can be derived from the analysis of DNA sequencing data alone. Slightly more formally, each process is characterized by a relative preference for each of the 192 types of all possible single nucleotide mutations in trinucleotide contexts oriented to the reference strand. Each process is assumed to vary along the genome and the observed heterogeneity of mutational spectra between loci is driven by different relative contributions of the processes (Fig. 1A). Inference of mutational processes then represents a classical blind source separation problem that separates a set of source signals from observed signal mixtures. For that, we devised a computational approach that performs dimensionality reduction using Principal Component Analysis (PCA) following by Independent Component Analysis (ICA) of mutational spectra in reduced space, so that processes have independent spectra and each process may have either positive (enrichment) or negative (depletion) preferences for context-specific mutation types (Fig. S1A, see Methods). Although there could be different mathematical formulations of a source separation problem, we argue that PCA-based dimensionality reduction following by ICA-based spatial inference is both statistically powerful and biologically reasonable for the population datasets considered here compared to the other state-of-the art approaches (Fig. S1H, see Methods). Simulations show that, accounting for the size and properties of the TOPMed dataset, our approach recovers processes that have a genome-wide contribution of at least 0.1% of the overall mutation rate and spatial scale of at least 10kb (Fig. 1E).

**Fig 1.**
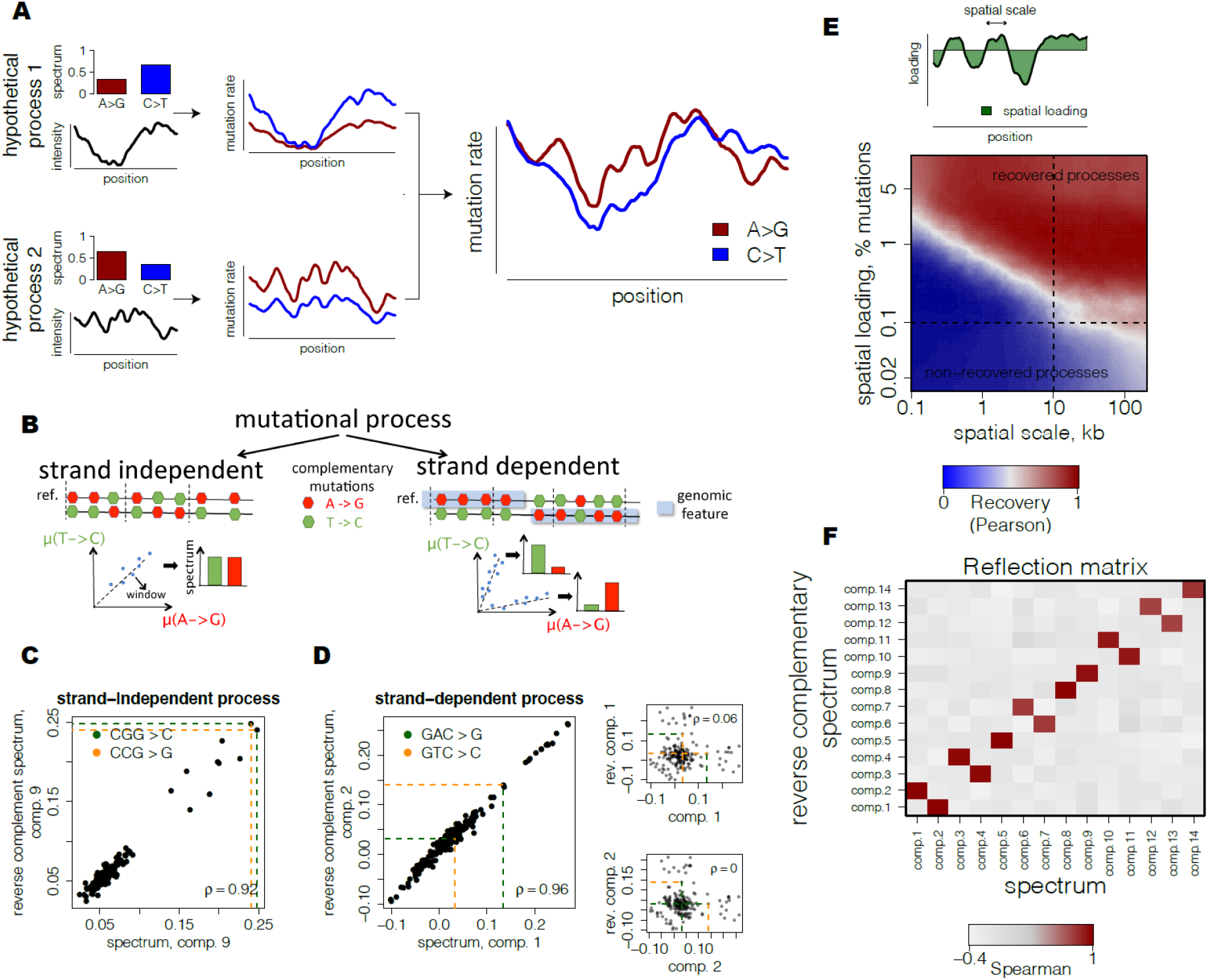
Inference of spatially-varying mutational processes in germline. **(A)** Observed spatial variability of mutational spectrum is modeled as a number of mutational processes with specific spectra and spatially-varying intensities. **(B)** Strand-independent mutational processes have equal rates of complementary mutations at each locus. Strand-dependent mutational processes produce two unequal patterns of complementary mutation rates at loci depending on the strand orientation of a genomic feature. **(C)** Example of a predicted strand-independent process. Loadings of complementary mutations of mutational component 9 are highly similar. **(D)** Example of a predicted strand-dependent process. Loadings of complementary mutations between two mutational components 1 and 2 are highly similar and characterize a single mutational process (left). In contrast to strand-independent process (see Fig. 1C), loadings of complementary mutations of mutational component 1 (upper right) and component 2 (lower right) are almost uncorrelated. **(E)** Theoretical scale-loading limitations for detection of mutational processes shows potentially high range of processes that can be recovered with the proposed approach. Simulations of mutational processes at different scales (quantified as half-life of simulated Ornstein-Uhlenbeck process) and spatial loadings (fraction of spatially-varying mutations of a process among total mutations, scheme) were based on parameters from TOPmed dataset. Quality of recovery was assessed using maximum absolute correlation between spectra of each simulated component and reconstructed components. **(F)** Reflection matrix reveals strand-dependency of processes and separates biological signal from noise. Correlation of spectrum of one mutational component with reverse complementary spectrum of another demonstrates clear separation into self-correlated components (5,8, 9,14) and pairs of mutually correlated components (1/2, 3/4, 6/7, 10/11, 12/13).

As with any statistical procedure, the key question is whether a particular inferred process reflects the biological reality or is a spurious signal. A powerful way to assess the biological relevance of the inferred processes is provided by the symmetry between antiparallel strands of DNA. Although DNA is a symmetric molecule, directional processes such as transcription and replication break this symmetry. Mutational mechanisms coupled with these processes are strand-dependent. For example, within genes A>G mutations are depleted on the transcribed strand and enriched on the complementary non-transcribed strand. This observation is attributed to the action of transcription-coupled repair (TCR) (*3*, *14*). All mutational mechanisms can be broadly classified into strand-dependent and strand-independent.

Our statistical procedure assigns the direction of mutations with respect to the human genome reference irrespectively of the direction of transcription, replication or double strand break repair. For some genes the reference strand happens to be transcribed, while for other genes it happens to be non-transcribed. As a consequence, in some genic regions we will detect depletion of A>G mutations and in others we will detect depletion of its complementary mutation T>C.

For a strand-dependent mutation process, our statistical procedure would infer two independent components (Fig. 1B). Remarkably, these components can be easily identified as corresponding to the same underlying process because they would be exactly complementary to each other. Following the example of mutation processes associated with transcription, the intensity of A>G mutations in one of the components would be identical to the intensity of T>C in the other. In contrast, a mutation process that is not strand-dependent would generate a component that would be self-complementary (for example, the intensity of A>G would be identical to the intensity of T>C). As a result, all biologically relevant components would either be self-complementary or arise in mutually complementary pairs (Fig. 1B-D).

We rely on this observation to test the biological validity of the inferred processes. Motivated by the visual representation in Figure 1B,F, we called this test a “reflection test”.

We applied our method to a dataset of very rare single nucleotide variants (SNVs) from the TOPMed freeze 5 (*15*) serving as a proxy to mutations (*16*). Overall, the dataset included over 293 million SNVs with allele frequency below 10^−4^. To capture the regional variation, we binned the genome into 264,291 non-overlapping windows of 10 kb, which is the optimal scale for the number of inferred components (Fig. S1G)

ICA identifies 14 independent components that successfully pass the “reflection test”, corresponding to 9 mutational processes, 5 of which are strand-dependent and the remaining 4 are strand-independent (Fig. 1F, Fig. S1C-E, Fig. S2). Almost all of these components have the average bootstrap support at the level of 70-99% (Fig. S1D).

These 14 components are robust with respect to window size and are reproduced in the independent gnomAD dataset (Fig. S1F). Finally, we validated these components using *de novo* mutations identified by parent-child trio sequencing (*17*, *18*). The spectra of *de novo* mutations in loci dominated by a specific component show a high concordance with the component spectrum inferred from the TOPMed dataset (Fig. S1K-M).

Eight of nine processes show notable and highly distinct correlations with genomic features known to impact mutation rate, including gene bodies, replication timing, direction of replication, and chromatin accessibility (Fig. 2A, Table S1). This strong association is remarkable given that the statistical inference was totally agnostic with respect to features other than mutation density.

**Fig 2.**
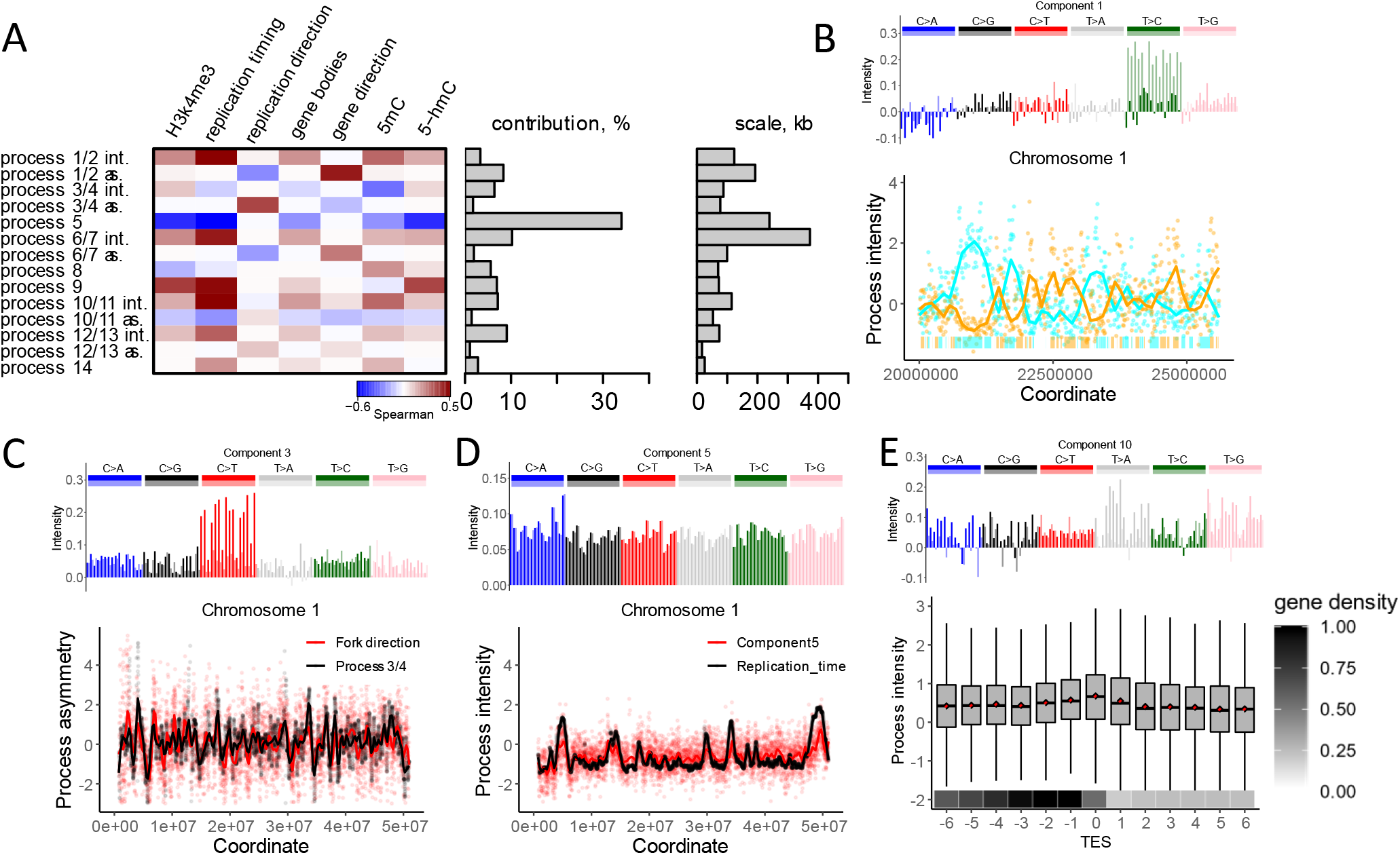
Mutational processes are associated with distinct genomic features. **(A)** Heatmap of correlations of intensities with genome features shows diverse modes of associations (left). For strand-dependent processes two spatial characteristics were considered: intensity (int.), estimated as the sum of intensities of two components, and asymmetry (as.), estimated as the difference between intensities of two components. Fraction of mutational variance explained by each process (middle) and scale, estimated as the half-life of the autoregressive model (right) are shown. **(B)** (top) The spectrum of one of the two components comprising process 1/2; (bottom) An example of intensities of components 1 and 2, associated with non-transcribed strands on chromosome 1. The bars on the bottom of the panel depict gene bodies (colors: cyan if transcribed strand is the reference strand and orange otherwise). **(C)** (top) The spectrum of one of the two components comprising process 3/4; (bottom) The association between the asymmetry of process 3/4 (component 3 – component 4) and the direction of the replication fork measured as a gradient of replication timing. **(D)** The spectrum of component 5 (top), and its association with replication timing (bottom). **(E)** (top) The spectrum of component 10; (bottom) Intensity of component 10 among 13 consecutive 10 KB-long regions adjacent to the transcription end site (TES). Box plot shows component 10 intensity. Mean intensity of component 10 in each region shown as a red point. Mean fraction of the transcribed nucleotides per region shown on the bottom.

Broadly, mutations can be introduced either as replication errors or as a consequence of DNA damage. The hallmark of mutations induced by bulky DNA damage is strand asymmetry with respect to direction of transcription (*3*, *19*) and, as we recently argued, direction of replication (*20*). Bulky DNA damage is resolved in a strand specific manner within gene bodies due to the action of TCR (*3*, *21*) and due to the preferential error-prone damage bypass on the lagging strand during replication (*2*). Components 1 and 2 have mutually complementary spectra and together correspond to a single strand-dependent process (Fig. 1D, Fig. 2A, Fig. S1). The strand asymmetry of this process, measured as the difference between intensities of components 1 and 2, strongly correlates with directions of both transcription (r=0.32) and replication (r=−0.15). The sum of the two components intensities reflects the overall regional activity of the process 1/2. For the process 1/2, it correlates with replication timing (r=0.34). Components 1 and 2 correlate in strand specific manner with the experimentally obtained activity of the transcription coupled repair system (*21*, *22*) in a strand-specific way (Fig. S3). Collectively, these observations strongly suggest that the process 1/2 is driven by the asymmetric resolution of bulky DNA damage.

In contrast, strand-dependent process 3/4 likely captures replication errors. The asymmetry of this process strongly correlates with the direction of replication (r=0.31) but is not meaningfully associated with any other epigenomic feature including direction of transcription. Therefore, in contrast to process 1/2, this process is unlikely to be mediated by bulky DNA damage. We hypothesize that process 3/4 reflects either a differential fidelity between replicative polymerases or a differential efficiency of mismatch repair (MMR) between leading and lagging strands (*1*, *23*, *24*). Although replication infidelity is frequently assumed to be a major (or even leading) factor in germline mutagenesis (*25*, *26*), process 3/4 offers the first probable genomic footprint of replicative errors. Interestingly, process 3/4 (sum of intensities of components 3 and 4) does not appreciably correlate with replication timing, even though many other processes do.

Process 5 most closely tracks replication timing (r=0.54), showing greater intensity in late-replicating regions. The association of germline mutation rate with replication timing was noted a decade ago, but it was shown to be quantitatively weak (*13*, *27*). A recent study reported that the association is much stronger for C>A mutations (*28*). C>A mutations are indeed enriched in process 5, although this enrichment is limited to TpCpN sequence contexts. Unlike other processes, process 5 affects all mutation types in the same direction (all types have positive values in the spectrum). This process is responsible for the largest fraction of mutation rate variation along the genome (Fig. 2A). In spite of these observations, the interpretation of this process is not straightforward because replication timing itself is correlated with many epigenomic features. Interestingly, most other processes are associated with replication timing, not only to a weaker degree, but also in the opposite direction (Fig. 2A). This counteracting effect explains the weakness of the association between overall mutation rate and replication timing.

Strand asymmetric process 6/7 is dominated by C>G transversions and is characterized by strong local spikes totaling 265 Mb throughout the genome (Figure 3A-C). Analysis of *de novo* mutations within these regions reveals that they are dramatically enriched in mutations of maternal origin (Table S2). Several genomic regions with high prevalence of maternal mutations, many of them occurring in clusters, have been reported by the original trio sequencing studies (*29*, *30*). Spikes of process 6/7 include all these regions and many previously unreported regions, also strongly enriched in individual and clustered mutations of maternal origin (Table S2 and Table S3). Overall, the rate of clustered maternal *de novo* mutations in regions of high intensity of process 6/7 is 18-fold higher than in the rest of the genome. These regions constitute 10% of the genome but harbor 67% of clustered maternal mutations (Fig. 3D, Table S3). Mutations in high intensity regions of process 6/7 have stronger dependence on maternal age and are responsible for 35% of mutations caused by oocyte aging. Mutations within these regions show a 2.6-fold excess in children of older mothers compared to younger mothers (Fig. 3H). In the remaining 90% of the genome this excess is just 1.4-fold. In contrast to earlier reports, this difference is not limited to C>G mutations (*30*).

**Fig 3.**
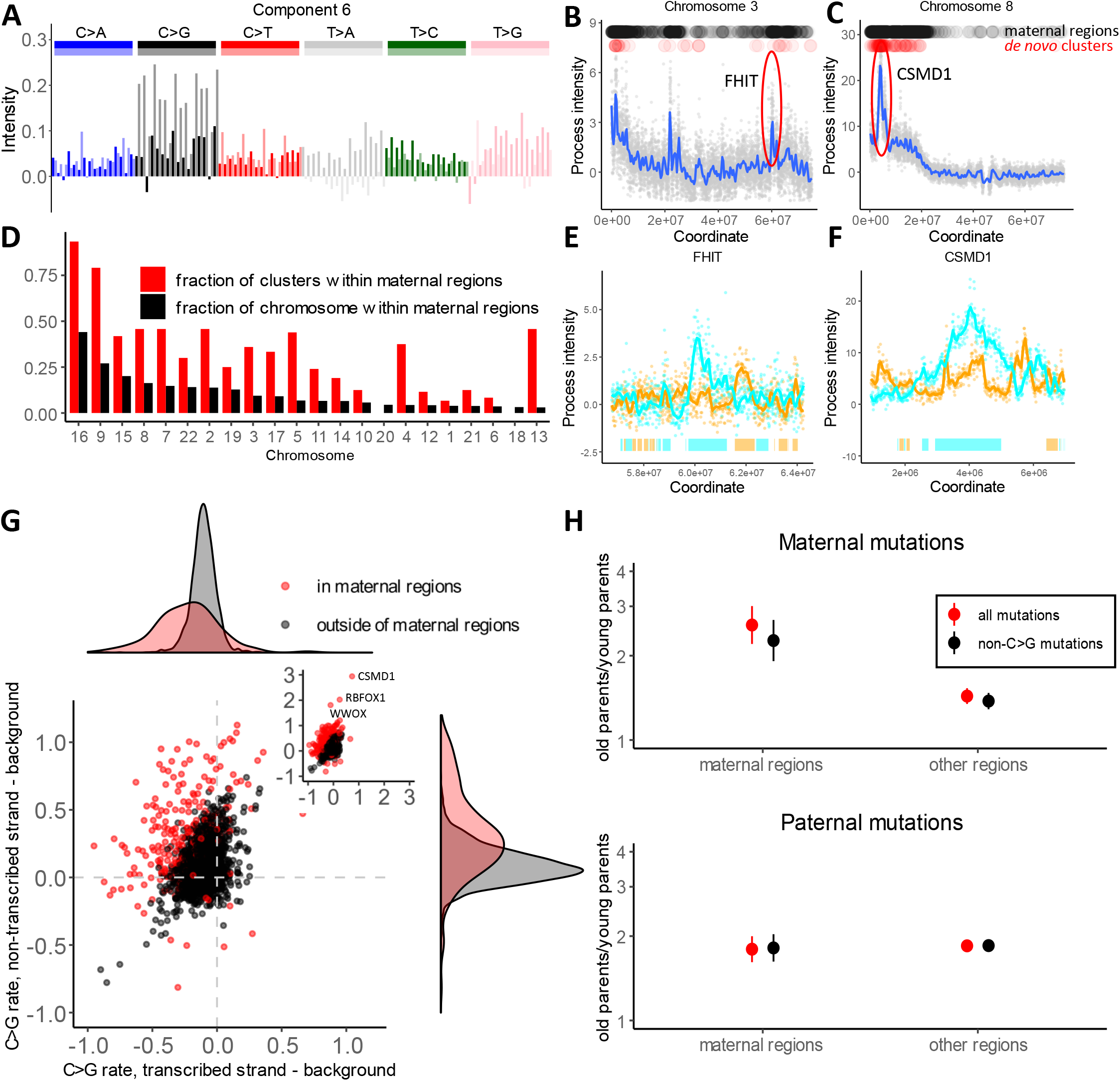
Oocyte-specific mutational process. **(A)** The spectrum of one of the two components comprising process 6/7. **(B and C)** Examples of two loci with high intensity of process 6/7, estimated as the sum of intensities of component 6 and component 7. Black dots on top of the panels mark windows of high intensity that we call “maternal regions” (see Methods). Red dots show *de novo* maternal clustered mutations from Halldorsson et al. (*18*). (**D**) Enrichment of maternal clustered de novo mutations from Halldorsson et al. in maternal regions. The fraction of each chromosome that is attributed to “maternal regions” (high intensity of process 6/7) is shown in black. The fraction of maternal clustered mutations located within maternal regions on each chromosome is shown in red; the difference in size between the red and black bars indicates enrichment of clustered mutations within “maternal regions”. (**E** and **F**) Zoom in view of process 6/7 intensity spikes around FHIT and CSMD1 genes on non-transcribed strands. Bars on the bottom depict gene bodies (colors: cyan if transcribed strand is the reference strand and orange otherwise). **(G)** Difference between C>G mutation rate on transcribed or non-transcribed strand of a gene compared to a 100 KB region flanking the gene. Red dots correspond to genes within maternal regions and black dots corresponds to genes outside of maternal regions. Density plots on the top and right summarize the distributions on the X and Y axes. **(H)** Ratio of parent-specific de novo mutation rates between first and last parent age quartiles is shown, estimated independently for “maternal regions” and for the rest of the genome. The error bars show the 95% confidence interval (95% CI) for the ratio of two binomial proportions test.

Five prominent spikes of process 6/7 overlap long fragile genes (*WWOX, RBFOX1, CSMD1, FHIT, SDK1*) (*31*). In these and other genes, process 6/7 displays a strong strand asymmetry with respect to transcription (Fig. 3, Fig S4, Fig S5; r=0.26). Within the gene bodies as compared to flanking regions, the rate of C>G mutations is decreased on the transcribed strand and is increased on the non-transcribed strand by as much as 50-200% (Fig. 3, Fig. S5).

Maternal mutations accumulate in oocytes that are arrested in the second phase of meiosis from the early stages of embryogenesis. Thus, the age-related increase of maternal mutations is unlikely to be explained by replication errors. Alternative mutation mechanisms should involve either DNA damage or resolution of double strand breaks outside of S-phase. The latter is favored by the current literature (*29*, *30*). This is an appealing explanation in light of mutation clusters and the striking maternal age dependency resembling the impact of age on structural variants (*32*). This is consistent with our observation that process 6/7 overlaps genes with common fragile sites. At the same time, the directly established spectrum of mutations induced by recombination has no sign of enrichment in C>G and is very different from process 6/7 (*18*). Furthermore, the signature of homology repair deficiency in cancer genomes also has a very different spectrum (*4*).

The strand asymmetry of process 6/7 cannot be easily explained by the double strand break model. The reduction of mutations on the transcribed strand suggests the role of bulky DNA damage repaired by TCR. In addition, the relationship with direction of replication (r=0.14, Fig. 2A, Fig. S4A) probably indicates that the unrepaired lesions on the leading and lagging strands are asymmetrically converted into mutations at the very first division of the zygote. The most surprising observation is the increase of mutation rate on the non-transcribed strand. Transcription-associated mutagenesis (TAM) has been previously reported in lower organisms and in some cancer types (*19*, *33*). Our analysis identifies TAM in human oocytes and shows that it is primarily localized to bursts of the process 6/7 (Fig. 3G, Fig. S5). TAM is a strand-dependent process associated with transcription and is unlikely to be explained by double strand break repair. Collectively, these observations, point to the localized susceptibility to DNA damage or the failure of DNA repair.

Processes 8 and 9 are dominated by mutations in the CpG context. Process 8 is characterized by CpG transitions and describes a well-known mechanism of spontaneous deamination of methylated cytosines which converts them into thymines. As expected, the intensity of process 8 is positively correlated with methylation levels and is low in CpG islands marking actively demethylated regulatory elements. Process 9 is characterized by CpG transversions. The intensity of this process spikes at CpG islands and is negatively correlated with methylation level (Fig. 4). CpG transversions were previously shown to be positively associated with the level of cytosine hydroxymethylation (*34*). Based on high intensity in CpG islands, the negative correlation with methylation level and the positive correlation with hydroxymethylation level, we hypothesize that process 9 is caused by active demethylation of regulatory regions. Enzymatic demethylation is initiated by oxidation of a methylcytosine resulting in a hydroxymethylcitosine (*35*). The hydroxymethylcitosine base, following cycles of subsequent oxidation, is removed by the Base Excision Repair system (BER), creating an abasic site. Unfinished repair of abasic sites is known to result in C>G mutations (*36*).

**Fig 4.**
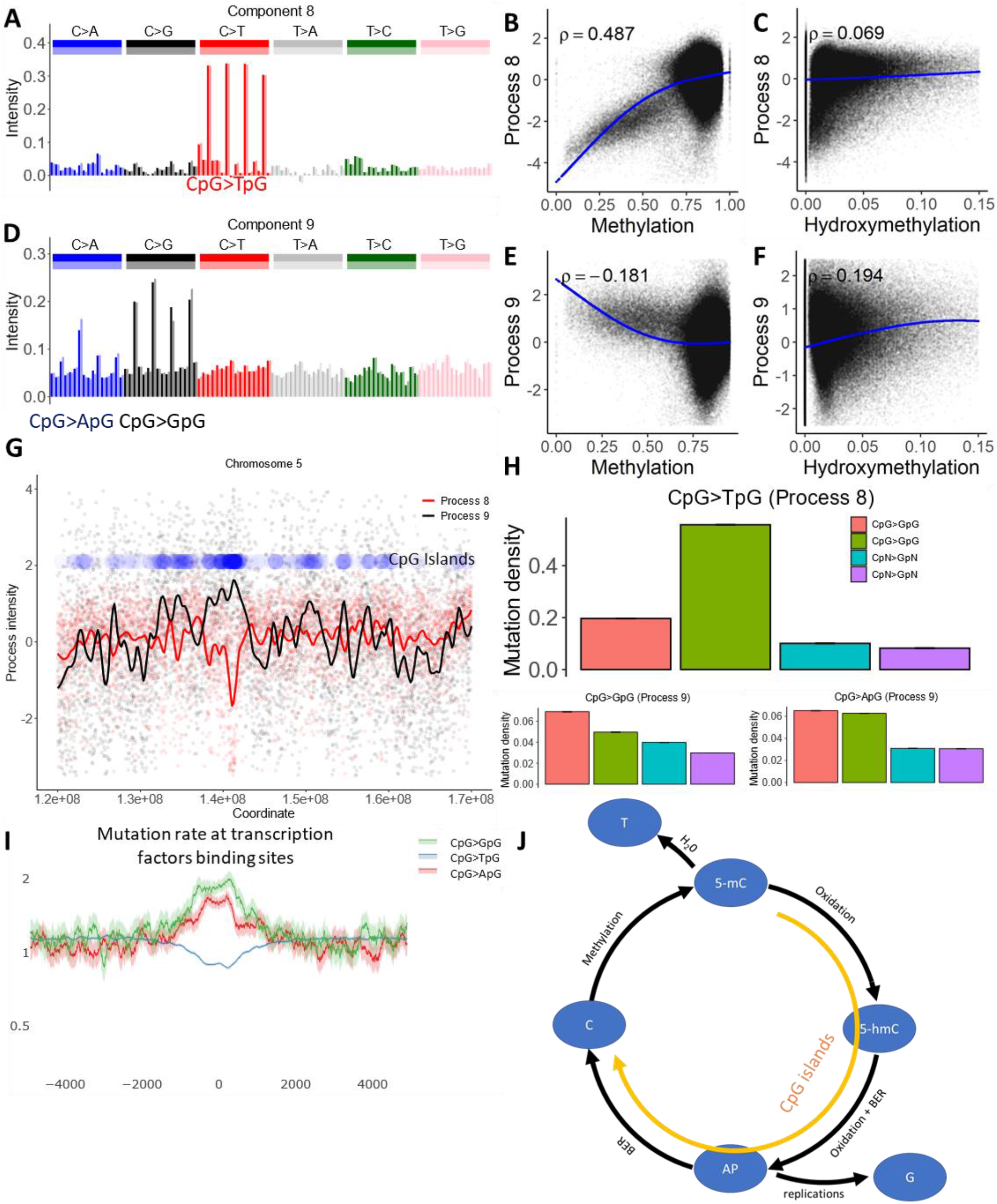
Cytosine deamination and cytosine demethylation. (**A** and **D**) The mutational spectrum of component 8 is dominated by CpG>TpG (top), while component 9 is dominated by CpG>GpG and CpG>ApG (bottom). (**B, C, E and F**) Association between the intensity of components 8 and 9 and cytosine methylation or cytosine hydroxymethylation. **(G)** Process 8 is inversely correlated with the density of CpG islands, while process 9 is positively correlated. Blue dots represent density of CpG islands across a 50 MB long region of chromosome 5. **(H)** Effect of CpG islands on mutation rate in CpG context and in cytosines outside of CpG context. **(I)** Mutation rate in CpG context at transcription factor binding sites located outside of annotated CpG islands, as determined by ChIP-seq peaks (see Methods), normalized to the genome average mutation rate. 95% binomial proportion confidence intervals are displayed in transparent lines. Higher levels of demethylation at these sites lead to the accelerated rate of CpG transversions. **(J)** Illustration of the biochemical mechanisms suggested for processes 8 and 9. Enzymatic oxidation of methylcytosine (5-mC) leads to hydroxymethylcytosine (5-hmC) (*35*), which after additional steps of oxidation should be removed by glycosylase, leaving an abasic site (AP). During DNA replication, AP sites will frequently be converted to CpG>GpG mutations and more rarely to CpG>ApG mutations, matching the spectra of process 9 (*36*). Alternatively, successful repair of AP sites creates non-methylated cytosines. Alternately, spontaneous deamination of methylcytosine creates a T to G mismatch, enhancing the rate of CpG>TpG mutations. While deamination should be prevalent in CpG sites with high methylation levels, the mutagenic effect of demethylation should be prominent in CpG islands.

Process 9 explains a small portion of the mutation rate variability. However, it disproportionately contributes to regulatory regions of the human genome. In undermethylated regions, the rate of CpG transversions is elevated under ChIP seq peaks for transcription factors (Figure 4). The mutagenic effect of repair of hydroxymethylated cytosines has been shown previously (*34*). We identify this process in an unsupervised manner and attribute it to unintended side effect of the functionally significant demethylation. In line with our model, cadmium, that suppresses cytosine demethylation, leads to depletion of C>G mutations in daphnia (Supplementary Manuscript).

The only remarkable association between intensity of process 10/11 and genomic features is a weak spike at the transcription end site on the transcribed strand of the gene (Figure 2E). Potentially this process is associated with transcription termination, but this localized effect is diluted at the 10 KB scale. The remaining processes 12/13 and 14 explain small proportions of the mutation rate variation. Statistical analysis of these processes does not unequivocally suggest specific biological mechanisms (see Supplementary Note for discussion of these processes).

Our analysis was enabled by the massive scale of the TOPMed dataset. Subsampling of the dataset shows that many components would not be detectable in smaller datasets. Even at the TOPMed scale, there are no statistical signs of saturation for the number of detectable processes (Fig. S1I-J) and a notable range of mutational processes remains undetectable in current settings (Fig. 1E). We hypothesize that larger population sequencing datasets are needed to paint a more detailed picture human germline mutagenesis.

In sum, our unsupervised statistical analysis of the genomic variation in mutation rate evident in population sequencing data implicates a compendium of biological processes responsible for human mutation. Our approach identifies a highly localized strand-dependent process dominated by mutations of maternal origin. This process tracks direction of transcription, suggesting a dominant role of transcriptionally-mediated damage in oocytes. We also characterize a mutation signature of replication errors, which has been historically suspected to be a major source of germline mutation. We attribute mutagenic patterns of repair of hydroxymethylated cytosines (*34*) to active demethylation of regulatory regions. We envision that a spatial mutational model applied to new datasets will uncover new links between DNA biochemistry and localized mutational patterns.

## Acknowledgements

Whole genome sequencing (WGS) for the Trans-Omics in Precision Medicine (TOPMed) program was supported by the National Heart, Lung and Blood Institute (NHLBI). Specific funding sources for each study and genomic center are given in Supplementary Note 3 and Table S4.

Centralized read mapping and genotype calling, along with variant quality metrics and filtering were provided by the TOPMed Informatics Research Center (3R01HL-117626-02S1; contract HHSN268201800002I). Phenotype harmonization, data management, sample-identity QC, and general study coordination were provided by the TOPMed Data Coordinating Center (3R01HL-120393-02S1; contract HHSN268201800001I). We gratefully acknowledge the studies and participants who provided biological samples and data for TOPMed. R.A.S and P.V.K. were supported by NHLBI (R01HL131768).

The views expressed in this manuscript are those of the authors and do not necessarily represent the views of the National Heart, Lung, and Blood Institute; the National Institutes of Health; or the U.S. Department of Health and Human Services.

## Material and Methods

### Preparation of mutational matrix

As a proxy for germline mutations, we used SNVs with allelic frequency below 10^−4^ from TOPMed freeze 5 (*1*) or gnomAD (*2*). The genome was binned into non-overlapping windows of 2, 5, 10, 30, 100 or 1000 kilobases in size, and mutation rate within each window was estimated as a ratio between the number of mutations and the number of available sites. To explore uniformity of the calling/sequencing quality, we obtained the distribution of the number of mutations within 1 kb windows across the genome. This distribution was bimodal with the first mode equal to 0 SNVs per region (Fig. S1A). This mode clearly corresponded to regions of low quality. Therefore, we excluded 1kb loci with the abnormally low mutation counts (less than 50 mutations) from all subsequent analyses. Overall, our results were stable with respect to different filtering thresholds (data not shown).

### Inference of mutational components

Mutation rates for each mutation type were Z-score transformed across all windows to zero mean and unit standard deviation. Using a predefined number of components ***n***, matrix *R* of transformed mutation rates in *w*=264’291 windows of *t*=192 mutation types was then factorized by singular value decomposition using the R package *svd*:

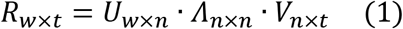

Matrix *V*_*n*×*t*_ of loadings of mutation types onto first ***n*** principal components was then centered to zero mean of columns *V*^_*n*,*t*_ = *V*_*n*,*t*_ − ⟨*V*⟩_*n*_, and rotated to infer statistically independent residual spectra components *M*^_*n*×*t*_ using the independent component analysis (ICA) R package *icafast*:

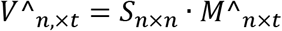

Components spectra were then defined as:

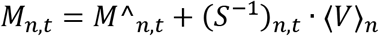

Since ICA defines components up to a sign and scalar, signs of rows of *M* were oriented to enable positive third moment and scales of rows were normalized to unit Euclidean norm. Oriented matrix *M*_*n*,*t*_ was considered as a matrix of normalized loadings of mutation types on components, while the matrix of intensities of mutational components in windows was estimated as: *I* = *U* · *Λ* · *S*. Altogether, matrix *R*_*w*,*t*_ of transformed mutation rates was factorized into a product of intensities *I*_*w*,*n*_ and independent spectra *M*_*n*,*t*_ of ***n*** mutational processes:*R*=*I* · *M*.

The spearman correlation coefficient was estimated between the spectrum and reverse complementary spectrum of each pair of components and with itself (we call it reflection correlation). Components having a reflection correlation more than 0.75 with at least one component were considered having reflection, or were otherwise considered to be noise. Inferred components were then classified into strand-independent, strand-dependent pairs or noise using the reflection test. Among components with reflection, components having a reflection correlation of more than 0.75 to itself were considered strand-independent. Pairs of components having a reflection correlation with each other were considered as two components of a strand-dependent process. Empirical observations show that a cutoff of 0.75 falls in a wide interval of values that deliver the same classification of components.

Since the reflection property of a component likely indicates its biological relevance, we used the number of components having reflection as a natural criterion to choose a predefined number of inferred components ***n***: the number of components with reflection was estimated for the range of values of ***n*** from 2 to 50, and the value of ***n*** corresponding to the highest number of components with reflection was selected. The procedure identified that the maximum number of mutational components with reflection is 14 for 10kb genomic windows.

As a first step *R*_*wxt*_ matrix was factorized on 14 components (n=14) with *svd.* Than to select the optimal window size, the algorithm was applied for a range of windows from 2 kb to 1 mb (2, 5, 10, 30, 100, 1000 kilobases) using these 14 input components. The number of components with reflection (Spearman correlation > 0.75) was estimated for each window size. Only window of 10 kb had all 14 components with reflection.

### Power analyses of the datasets

The dataset was subsampled up to the size of 1, 5, 20, 40, 60, 80, 90 and 95 % of the original dataset. For each subsampled dataset the method estimated mutational components using *svd* decompose matrix as input. Recovery quality of a component of the original dataset in each subsampled dataset was estimated as maximum absolute Pearson correlation across all inferred components of a subsampled dataset. To account for uncertainty in subsampling outcome, quality of recovery was averaged across 10 independent sampling runs at each subsampling depth. Finally, at each subsampling depth we estimated average number of components 1) having reflection and 2) having highly correlated (>0.75) counterpart in the original dataset.

### Comparison of different inference methods

We compared methods that use decorrelation, independence, and non-negativity as constraints on matrix factorization problem. PCA was used as a baseline method that decorrelates mutational components using matrix *R* of mutation type rates, Z-score transformed across genomic windows. In case of PCA, mutational components were interpreted as rows of orthogonal matrix *V* (see equation 1). PCA was also used as a dimensionality reduction approach before ICA. Mutational components that maximize independence of spectra were obtained as described above, while independence of intensities was achieved using ICA of orthogonal matrix *U* (see equation 1).

On the other hand, non-negative matrix factorization (NMF) approach was applied to the matrix of mutation type rates with each mutation type rate normalized by its genome-wide average level. Using *NMF* R package we run standard NMF algorithm (option ‘brunet’), NMF that tends to produce sparser components (option ‘ns-NMF’, default parameters) and NMF that tends to diversify expression of components patterns (option ‘pe-NMF’, parameters: alpha=0.01, beta=1). Since we noticed that for this TOPMed dataset NMF tends to converge to different local optima, each NMF algorithm was run using 10 starting points, including ‘nnsvd’, ‘ica’ and 8 ‘random’ options. To make analysis of NMF components compatible with that of PCA and ICA, NMF-inferred components were centered by subtracting 1 (normalized genome-wide average rates). All of the methods were run using dimensionality of 14 of input components. Components that have spectra dominated by a single outlier mutation type, that is 10 times exceeding loadings of any other mutation types, were removed. Reflection test with cutoff of 0.75 on reflection correlation was used to estimate the number of potential biological components for each method.

### Statistical properties of mutational components

The scale of mutational components was defined using a linear autoregressive model. The spatial intensity of each mutational component was modeled as:

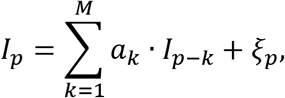

where *I*_*p*_ is the intensity at position p, *a*_*k*_ are autoregressive coefficients and ξ_*p*_ is the residual noise. Order *M* of the model was chosen using Akaike Information Criterion. The R package *ar* was used to fit the autoregressive model. The scale of each process was defined as the half-life of the autoregressive model

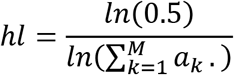

The contribution of each component was defined as the squared sum of intensities. Contributions of all components were then scaled to the unit sum.

### Assessment of components robustness

Robustness of each component spectrum was assessed using a bootstrap of genomic windows. 500 sets of 14 components were inferred using a bootstrap of windows. Maximum Spearman correlations between an original component and the components in each bootstrapped set were calculated to provide estimates of the similarity of potentially identical components. For all mutational components, average Spearman correlations of the spectra with bootstrapped components were above 0.68, indicating the robustness of spectra estimates.

Inference of components was repeated for a window size of 5 kb and 30 kb to explore robustness with respect to window size. For each window choice, the procedure of inference was repeated independently, including selection of the optimal number of components. The spectra of all original components were recapitulated, with a correlation of more than 0.64 in at least one of two runs. Finally, component spectra were compared between TOPMed and gnomAD datasets. For that, the procedure of components inference, including selection of the optimal number of components, was repeated for the gnomAD dataset using a window size of 100 kb. The spectra of most components were recapitulated with a correlation of more than 0.6, while three components (3, 10, 11) showed moderate correlation (0.46, 0.43, 0.55). Overall, this indicates that components are robust with respect to the choice of window and dataset.

### Comparison with *de novo* data

To assess if the spatial distribution of *de novo* mutations is consistent with individual mutational processes, we pooled 421,106 *de novo* point mutations from two datasets (*3*, *4*) and estimated the log ratio of *de novo* frequencies of mutation types in 25% of genomic windows of high component intensities relative to frequencies of mutation types in the whole genome. Consistency of *de novo* data with the mutational component is quantified as the Spearman correlation between this log ratio for *de novo* mutations and the spectrum of the corresponding mutational component. Spearman correlations were positive for each component. To estimate the uncertainty of these correlations, we repeated the estimation of Spearman correlations multiple times using bootstrapped sets of genomic windows. The significance of each association was assessed as the p-value of zero correlation relative to the distribution of bootstrapped Spearman correlations. The results show that for all components the correlation is significantly consistent (p < 0.05; Supplementary Figure 1). To assess parent-specific effects, the Spearman correlation between the log ratio of *de novo* mutation frequencies and the spectrum of mutational process was estimated separately for phased maternal and paternal *de novo* mutations. Before this procedure, 63,387 paternal *de novo* mutations were downsampled to match the size of 17,406 maternal de novo mutations. The distribution of differences between maternal and paternal Spearman correlations was constructed using a bootstrap of genomic windows to assess statistical significance, estimated as the p-value of zero correlation relative to the bootstrapped distribution. Similarly, to assess age effects of mutational processes, the dataset was partitioned by the average age of parents in two equal parts of young and old parents and the procedure identical to that applied for parent-specific effects was repeated.

### Simulations to assess the limitations of the approach

The ability to infer spatially-varying mutational processes depends on their statistical properties, such as spatial scale, degree of variability along the genome and degeneracy of mutational spectrum. Limitations of inference with respect to these statistical properties were analyzed through simulations of mutational processes underlying spatially variable mutation rates. Briefly, we simulated spectra and intensities of 14 mutational components corresponding to 4 strand-independent and 5 strand-dependent processes (4·1 and 5·2 components respectively), linearly combined them to obtain variable mutation type rates along the genome and sampled mutation counts using Poisson process. Then the procedure of spatial inference was made for the matrix of simulated mutation counts and spectra of inferred components were compared to simulated ones to estimate quality of recovery. Recovery quality of a simulated component was calculated as a maximum absolute Pearson correlation to inferred components. Simulations were repeated 8000 times to assess processes in a wide range of scales, loadings and spectra degeneracies. In more detail, intensities of components were simulated by continuous Ornstein-Uhlenbeck (O-U) processes. Scale of component was estimated as half-life (*hl*) of O-U process. The latter was sampled from 100 bp to 200 kb uniformly at log scale. Stationary O-U mean *m* (see equation 2) was assigned to 3 and stationary variance *α* was sampled from 0.005 to 5 uniformly at log scale. Stationary variance controls the degree of spatial variability of components. Overall, intensities *I* were modeled using O-U diffusion:

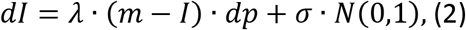

where p is genomic position, *λ is a rate of reversion* 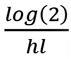,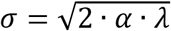,*α* = *exp*(*unif*(*log*(5 · 10^−3^),*log*(5))),*m*=3.

Rate vector of 192 mutation types was sampled using Dirichlet distribution 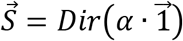 with a concentration parameter *α* sampled uniformly from 0.01 to 10 at log scale. Concentration parameter controls degeneracy of spectra and is shown in Supplementary Fig. 1a as a “spectra degeneracy” score. Mutation type rates of components spectra were then re-normalized to match the average observed genome-wide mutation frequencies in TOPMed. Rates *S*_*i*,*j*_ of each spectra mutation type *j* were scaled by a factor *c*_*j*_: *S*_*i*,*j*_ ← *S*_*i*,*j*_ · *c*_*j*_, where 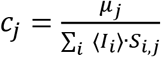 with *μ*_*j*_ being average genome-wide mutation rate of type *j* and ⟨*I*_*i*_⟩ is average intensity of a process *i*. Finally, expected mutation rates of each type *j* in each window *w* is a linear combination of components *v*_*w*,*j*_ = ∑_*i*_ *I*_*wi*_ · *S*_*ij*_. Mutation counts of each type in each window were sampled from Poisson process with a rate *m*_*w*,*j*_ = *v*_*w*,*j*_ · *c*_*j*_ proportional to mutation rate *v*_*w*,*j*_and average number of available *c*_*j*_ context triplets per window in the human genome. The procedure of components inference was then applied to matrix *m*_*w*,*j*_ of simulated mutation counts.

### Associations with epigenetic tracks and DNA features

We relied on the analysis of correlations between mutational processes and epigenomic tracks to gain insight into biological mechanisms.

Replication timing was obtained from (*5*). In the absence of data from the relevant germline tissue, we used the track for Mcf7 cells. The results were insensitive to the choice of cell type. Replication fork direction was determined as in (*6*).

Gene coordinates were obtained from the ‘knownGenes’ track downloaded from the UCSC genome browser. We measured gene bias within each window as the number of nucleotides transcribed on the reference strand minus the number of nucleotides transcribed on the strand complementary to the reference. Correlations with process 6/7 asymmetry, estimated as the difference in intensities of components 6 and 7, were calculated only in regions of high intensity of process 6/7 (component 6 + component 7 intensity >1.4).

Methylation level for each CpG dinucleotide were obtained from (*7*) and the methylation level of a window was calculated as a mean across all CpG sites within it.

Hydroxymethylation data was obtained from (*8*). Because this track is very sparse, similarly to previous study (*9*), we considered any CpG site with the fraction of hydroxymethylated reads exceeding 0.1 as hydroxymethylated. The hydroxymethylation level of a window was calculated as the fraction of hydroxymethylated CpG dinucleotides among all CpG dinucleotides.

Histone modifications H3k4me3, H3k27ac and H3k4me1 were downloaded from the UCSC genome browser. These tracks were obtained for human embryonic stem cells as a potentially relevant cell type.

Sex-specific recombination rate were obtained from (*3*).

CpG islands coordinates were downloaded from UCSC genome browser. Correlations between all tracks and mutational processes are listed in Supl. Table 1.

### Associations with the activity of nucleotide excision repair

Nucleotide excision repair (NER) effectively removes bulky lesions and its activity is partly governed by chromatin structure (*10*). Kinetics of CPD and 6-4PP repair by NER was measured in (*11*). Repair of 6-4PP occurs within less than an hour and thus is unlikely to be relevant for the mutagenesis that operates in the germline, because divisions of spermatogonia take many days and the dictate phase of oogenesis lasts for many years. Therefore, we focused on the repair of CPDs, a much slower process (*11*). The majority of UV-induced lesions occur in TT dinucleotides due to properties of UV radiation. To account for this bias, we normalized NER activity to TT dinucleotide content. Following this, we correlated NER efficiency with the intensity of each mutational process.

On the other hand, local activity of NER should be inverse to the amount of the damage that remains in DNA after 48 hours past UV-irradiation. We correlated mutational processes with the amount of unrepaired CPD damage (*12*), normalized to TT dinucleotide content. Correlations between NER activity and mutational processes are shown in Fig. S4.

### Clustered de novo mutations

In line with previous studies, we defined clustered *de novo* mutations as pairs of mutations observed in the same individual at distances less than 20,000 nucleotides (*13*, *14*). *De novo* mutations were obtained from (*3*) and entire clusters were attributed to be of maternal or paternal origin if there was at least one phased mutation of this origin. Clusters that have mutations on both the paternal and maternal haplotype were excluded.

### Alteration of mutation rate in gene bodies

To directly estimate the effect of transcription on mutation rate, we compare the mutation rate for each of 12 mutation types on the non-transcribed strand of the gene to the mutation rate 100 KB upstream and downstream of the gene (Fig. 3G and Suppl. Fig. S5). To reliably estimate the intensity of the process and the mutation rate within genes, only genes longer than 100 KB were considered. Differences in mutation rate between the gene and flanking region were normalized to the genome-average mutation rate for each corresponding mutation type.

### Maternal age effect in regions susceptible to process 6/7

“Maternal regions” were determined by the high intensity of component 6 + component 7. To choose the threshold for this sum, we compared quintiles of the distribution of component 6 + component 7 to that of the normal distribution and the deviated right tail of 8601 windows was used to define “maternal regions” (Fig. S.5).

To calculate the effect of maternal age, Broyden–Fletcher–Goldfarb–Shanno maximum likelihood algorithm was used (R package bbmle). We deal with the uncertainty contributed by non-phased mutations as in (*15*).

### Effect of transcription binding sites on mutation rate

Aggregate ChIP-seq peaks were obtained from ReMap2018 (*16*). CpG islands were excluded from the following analysis.

Mutation rate for the set of transcription factor binding sites was calculated in overlapping 100 nucleotide-long sliding windows for each trinucleotide context, then this rate was normalized on genome average mutation rate values and combined into the stated categories using a weighted average.

## Supplementary Figures Legends

**Figure S1.**
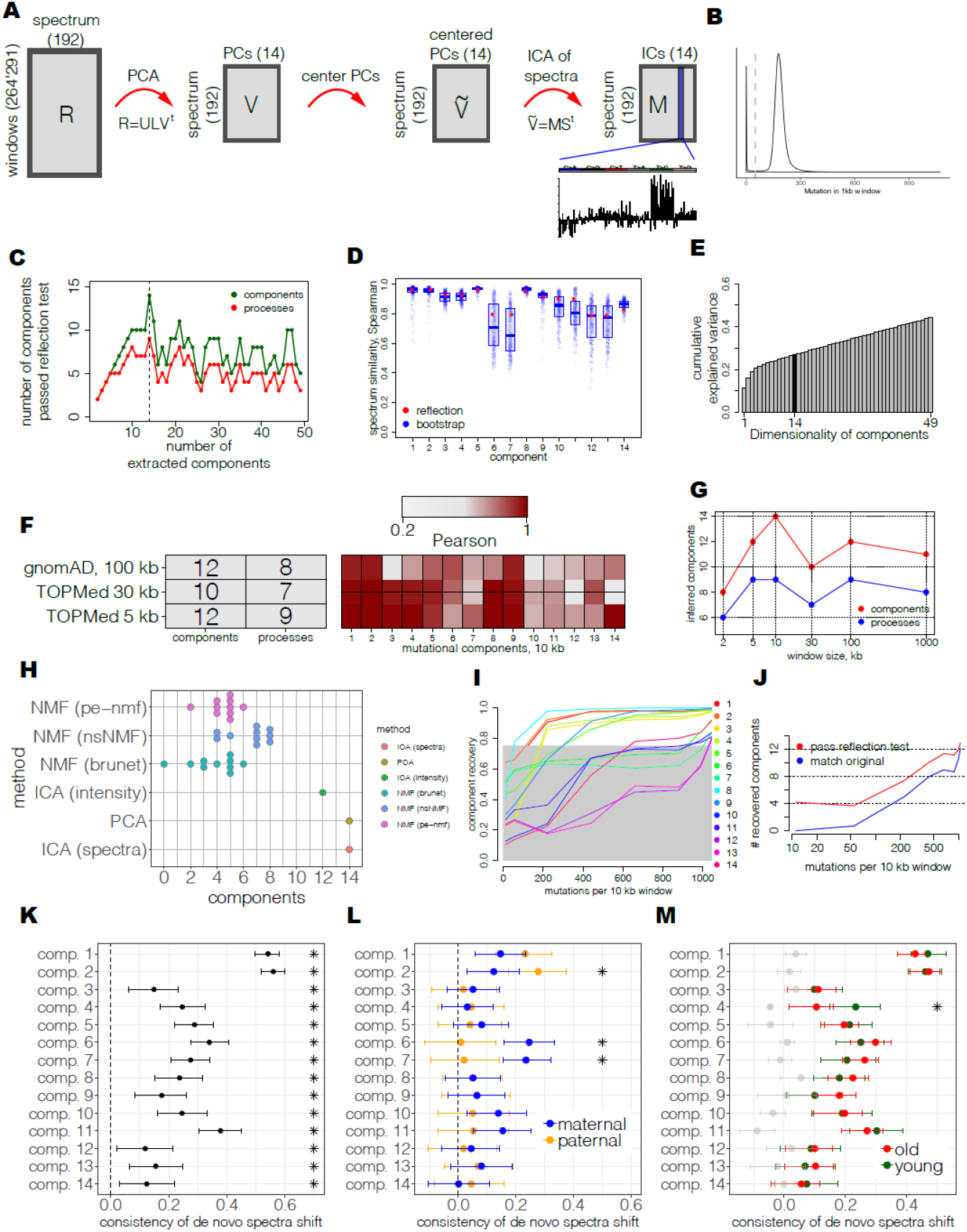
Statistical properties of inferred mutational processes. A) Computational approach. Matrix of mutation type rates (R), normalized to zero mean and unit variation across genomic windows, is reduced to 14 principal components to capture major axis of spatial mutational variability. Then, matrix of principal components is re-centered and rotated to maximize independence of spectra using ICA. B) Bimodal distribution of the mutation counts in 1 kb regions. Windows having less than 50 mutations were filtered out (grey bar). C) Reflection test provides a strategy to choose an optimal number of components. The number of components having reflection (spectrum similar to reverse complementary spectrum of any component among all extracted components) was calculated for a range of input numbers of components. The maximum number of components with reflection is 14. Quality of component spectra estimates through bootstrap and reflection property. For bootstrap strategy, similarity of extracted components with components extracted using bootstraps of genomic widows was assessed. A total of 500 bootstrap samples were generated. For each component, the most similar component in bootstrap sample was recorded to estimate confidence intervals (blue points). Alternatively, similarity of each extracted component spectrum to any reverse complementary component spectrum was assessed to estimate precision of spectrum (red points). D) Cumulative variance explained by extracted components. Black bar shows variance explained by 14 components used to predict mutational processes. E) Comparison of components spectra reproducibility of TOPMed with 10 kb windows, TOPMed with 30 kb windows, TOPMed with 5 kb windows and gnomAD with 100 kb windows. F) Assessment of performance using a range of windows shows optimality of 10 kb scale. The umber of components having reflection (inferred components) using 14 input principal components was estimated for windows of 2, 5, 10, 30, 100 and 1000 kilobases. G) Comparison of different methods shows superior performance of ICA of spectra and PCA. A number of components having reflection among 14 input dimensions (X axis) was compared for ICA of spectra, PCA, ICA of intensities and three implementations of non-negative matrix factorization (NMF): standard NMF (‘brunet’), non-smooth NMF (‘nsNMF’) and pattern-expression NMF (‘pe-NMF’). To account for the NMF property to converge to one of many local optima, 10 points per each NMF algorithm were generated to reflect results of different starting values (including ‘random’, ‘ica’ and ‘nndsvd’ options in NMF R package). Components that have strong outliers in spectrum were excluded (see Methods for details). H) Estimates of the dataset size sufficient to detect individual components. The original dataset was subsampled from 95% up to 1% (X axis) to estimate quality of recovery of each component (Y axis), measured as a maximum absolute Pearson correlation of an original component with components inferred from a subsampled dataset. Grey rectangle indicates area where components are not considered to be recovered. I) Number of detected components does not show saturation with increased size of the dataset. A number of components having reflection (red) or having highly similar components in the original dataset (blue) were estimated for a range of dataset subsampling depth (X axis). J) Spatial distribution of *de novo* mutations is consistent with intensities of mutational components. Comparison of spectra log ratio of 421,106 *de novo* mutations between regions of high and low intensity of a component and mutational spectrum of that component demonstrates correlated changes for all components. *De novo* mutations were pooled from Halldorssonet et al. (*1*) and An et al. (*2*) datasets. Mean correlation and 95% confidence intervals were estimated using bootstrap of genomic windows. Asterisks indicate p-value < 0.05, calculated as empirical p-value of zero correlation relative to distribution of 1000 bootstrapped correlations (see Methods for details). K) Parental-specific biases of mutational processes are detectable only for process 6/7. Consistency of log ratio shift in *de novo* spectrum with mutational component spectrum was estimated separately for phased maternal and paternal *de novo* mutations. Significance of parent-specific shift was estimated using empirical distribution of differential shift between maternal and paternal bootstrapped correlations among 1000 bootstraps. P-value was estimated as p-value of zero correlation with respect to constructed empirical distribution and asterisks show cutoff 0.05 on significance. L) Age-specific biases of mutational processes are detectable only for process 6/7. Consistency of log ratio shift in *de novo* spectrum with mutational component spectrum was estimated for top and bottom 50% of the dataset partitioned by the average parental age at conception. Significance of age-specific shift was estimated using empirical distribution of differential shift between older and young bootstrapped correlations among 1000 bootstraps.

**Supplementary Figure 2.**
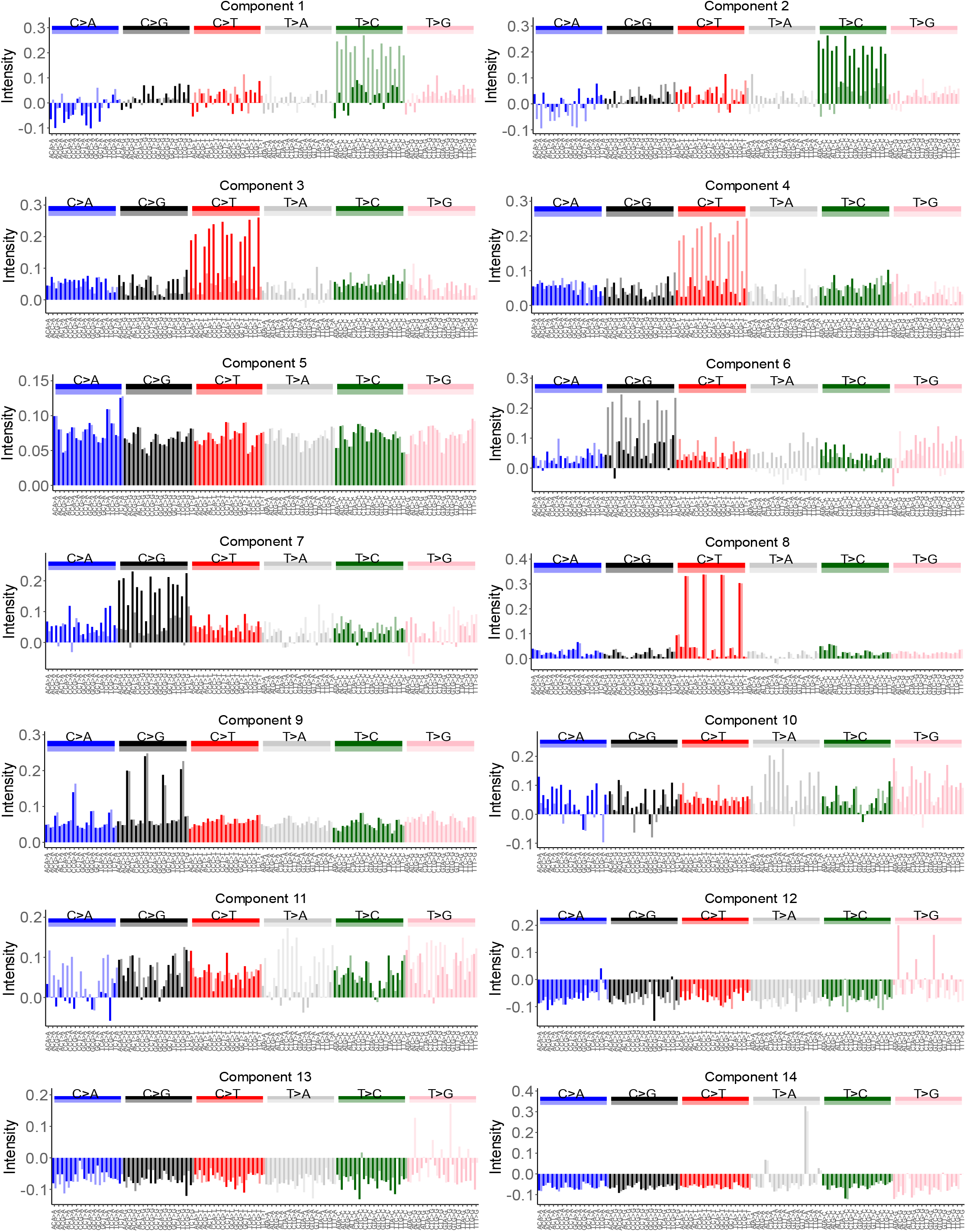
Mutational spectra of extracted components.

**Supplementary Figure 3.**
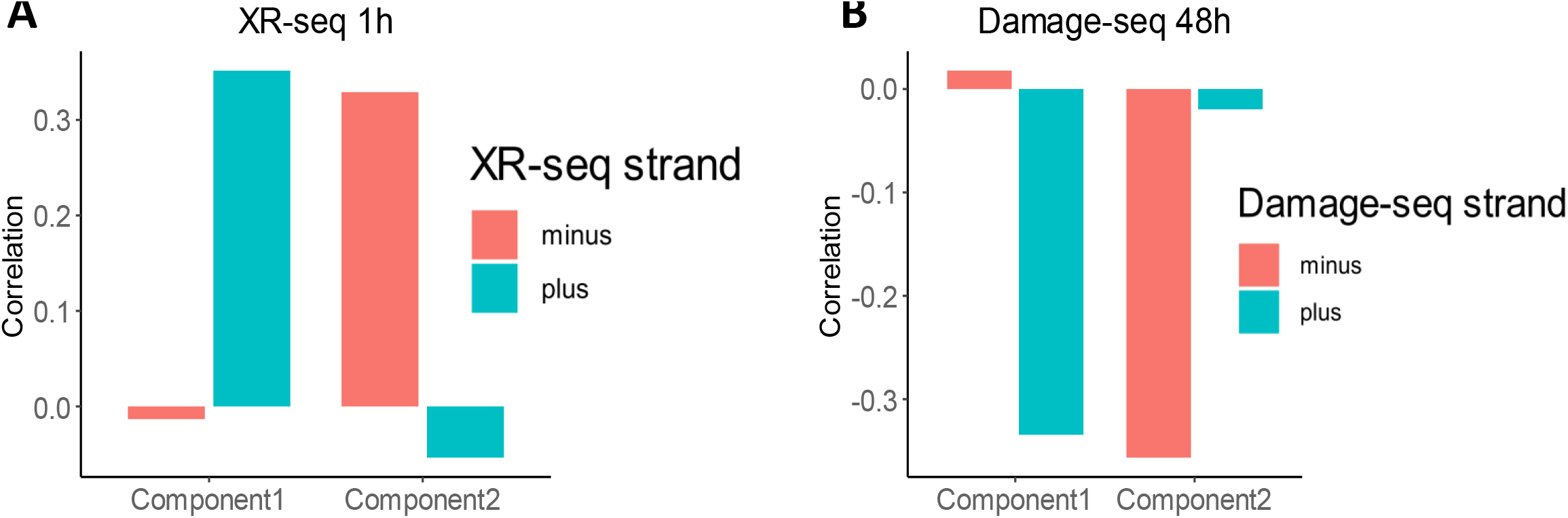
Processes 1/2 associated with experimentally measured NER activity. (A-B) Components 1, 2 show high correlation with (A) XR-Seq and (B) Damage-seq from (*3*, *4*) in a strand-specific way.

**Supplementary Figure 4.**
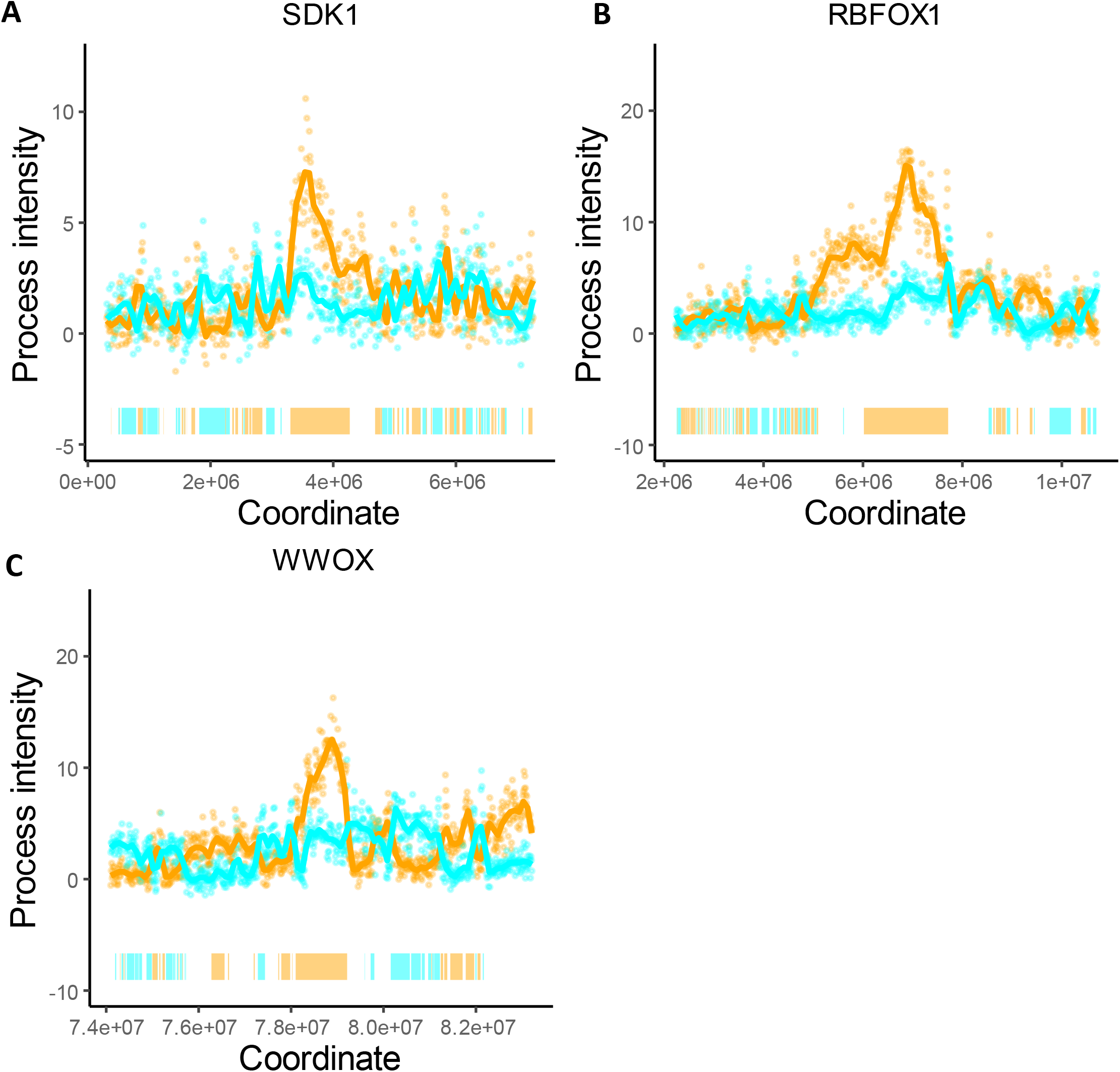
Spikes of the process 6/7. Process 6/7 intensity spikes around SDK1, WWOX and RBFOX1 genes on non-transcribed strands. Bars on the bottom depict gene bodies (colors: cyan if transcribed strand is the reference strand and orange otherwise).

**Supplementary Figure 5.**
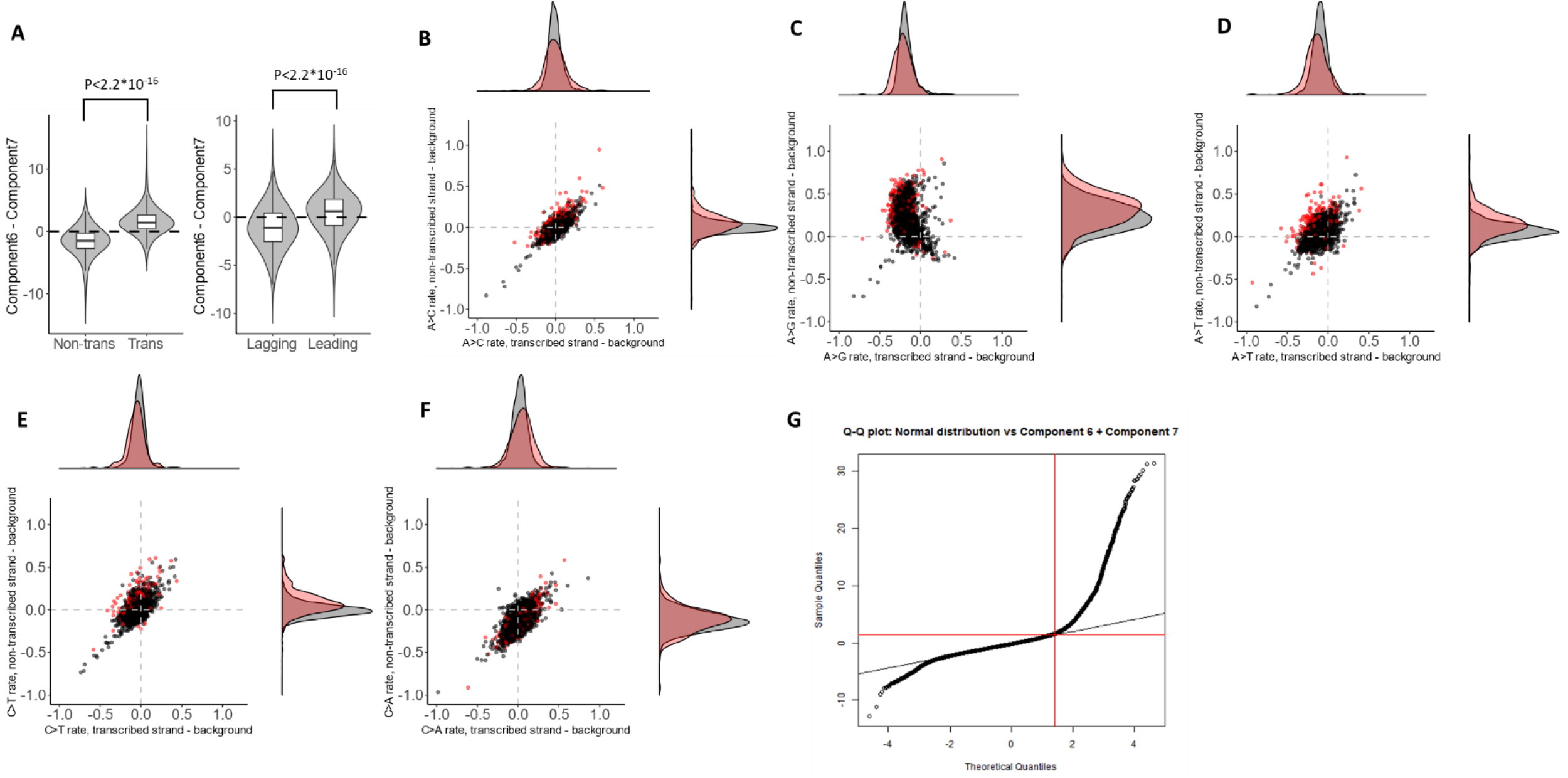
Properties of process 6/7. (A) Violin plots shows asymmetry of process 6/7 with respect to transcription and replication. The sign of asymmetry matches the direction of transcription in 81% of regions and matches the direction of replication in 64% of intergenic regions. (B-F) Mutation type-specific comparison of mutation rates on the transcribed and non-transcribed strand of a gene relative to 100 KB flanking regions. Red dots correspond to the genes within maternal regions and black dots correspond to genes outside of maternal regions. Density plots on the right and on the top summarize the distribution on Y and X axes. (G) We define regions with a high intensity of maternal signature as the left tail deviated from normality. Red line shows the chosen threshold equal to 1.4.

